# *In situ* three-dimensional mapping of oxygen gradients in *Staphylococcus epidermidis* biofilms using a solution-based, ratiometric imaging platform

**DOI:** 10.64898/2026.05.28.728557

**Authors:** Patryck J. Michalik, Elizabeth J. Stewart

## Abstract

*Staphylococcus epidermidis* biofilm oxygen gradients are spatially mapped during biofilm development and after vancomycin treatment using a solution-based ratiometric imaging platform. By integrating the oxygen-sensitive, tris(2,2’-bipyridyl)dichlororuthenium(II) hexahydrate with oxygen-insensitive, water-soluble CdSe/ZnS quantum dots, we achieve *in situ* microscale resolution of dissolved oxygen (DO) concentrations within biofilms. The oxygen-sensing platform is calibrated within alginate hydrogels to mimic probe confinement within biofilms and subsequently validated in biofilms using chemical oxygen depletion (sodium sulfite) and thermal inactivation (60°C). We demonstrate that the probes do not significantly alter planktonic bacterial growth or biofilm development. Using confocal laser scanning microscopy and quantitative image analysis, 3D microscale oxygen maps of biofilms are visualized and evaluated. During *S. epidermidis* biofilm development from 12 to 24 hours, average biofilm DO concentrations decrease from 2.62±0.22 mg/L to 2.02±0.47 mg/L, corresponding with increased *S. epidermidis* biofilm biomass and bacterial metabolic activity. While treatment of *S. epidermidis* biofilms with vancomycin at the minimum inhibitory concentration (MIC) (2 µg/mL) results in negligible DO decreases and biofilm biomass and metabolic activity comparable to untreated biofilms, higher vancomycin concentrations (20, 200 µg/mL) lead to increases in biofilm DO, higher dead-cell biovolumes, and decreased metabolic activity. This work establishes a microscale, solution-based ratiometric platform for quantifying the interplay between biofilm oxygen gradients, structure, and metabolic activity, providing a framework for understanding biofilm resilience during antimicrobial treatment.

## 1. Introduction

Bacterial biofilms are structured communities of cells encapsulated in a self-produced extracellular polymeric matrix^1^. Biofilms are ubiquitous in both natural and clinical environments^1^, where they can exhibit structural, chemical, and physiological heterogeneity^2, 3^. The structural and chemical properties of bacterial biofilms can vary dynamically in response to shifts in environmental conditions^4^. Throughout the lifecycle of the biofilm, fluctuations in the growth microenvironment and bacterial metabolism can lead to the formation of oxygen gradients within biofilms^2, 3^. As oxygen availability in the biofilm microenvironment varies, bacteria within biofilms adapt to the aerobic and anaerobic conditions within their surroundings^2, 3^. Generally, bacteria located deeper within the biofilm experience lower oxygen availability and differing metabolic activity compared to bacteria at the upper surface of the biofilm^2, 3^. Reduced oxygen availability can lead to regions of biofilms with low metabolic activity, which contributes to their recalcitrance to antibiotic treatment^5^. While oxygen concentration is an important indicator of bacterial metabolic activity and growth within biofilms, experimental tools to quantify oxygen gradients within biofilms at spatial and temporal scales relevant to bacterial metabolic activity remain limited^6^.

Oxygen gradients in biofilms have been measured using a range of techniques, including microelectrodes, two-dimensional planar optodes, and fluorescent nano/micro-particles^6^. Utilization of microelectrodes has allowed for high sensitivity and dynamic measurement of oxygen concentrations in bacterial biofilms^6^; however, microelectrodes are limited to point measurements and can perturb biofilm structure^6^. Two-dimensional planar optodes have been applied to measure biofilm oxygen concentrations at the substrate surface where biofilms grow, but do not spatially resolve oxygen concentrations within the three-dimensional structures of biofilms^6^. Nano/micro-particle based oxygen monitoring has been adapted for measuring biofilm oxygen concentrations in three dimensions^6^; however, current nano/micro-particle-based oxygen monitoring systems have diffusional limitations due to nano/micro-particle size, which generally ranges between 200-1000 nm^7–9^. Additionally, as bacteria are similar in size to probes between 200-1000 nm, the use of these probes may physically alter biofilm structures or oxygen gradients. Non-destructive, three-dimensional mapping of oxygen gradients in biofilms is important for advancing the understanding of the spatial heterogeneity of biofilm metabolic activity and response to antimicrobial treatment.

Ratiometric luminescent molecular probes have been utilized for oxygen imaging in living cells and tissues^10^. Ratiometric oxygen sensing is typically achieved by coupling an oxygen-sensitive luminophore (e.g. ruthenium (II) complexes) with an oxygen insensitive reference luminophore (e.g. quantum dots) to detect oxygen concentration^11^. Oxygen-sensitive ruthenium (II) complexes are frequently used for ratiometrically measuring oxygen^10, 11^. Ruthenium (II) complexes have been utilized in a planar coverslip-based O_2_ sensor to measure 2D biofilm oxygen concentrations^12^ and in 1 µm oxygen-sensing microparticles to measure 3D biofilm oxygen concentrations^7^. Quantum dots offer many advantages as oxygen-insensitive materials for ratiometric oxygen sensing due to their high photostability, narrow emission band, and high quantum yield and are frequently utilized as the oxygen-insensitive reference probe in oxygen sensing^13, 14^. An additional advantage of ruthenium (II) complexes and quantum dots for solution-based mapping of oxygen concentrations in biological samples is their size (<10 nm) as these complexes do not face the diffusion limitations of the larger nano- and microparticle sensing platforms.

*Staphylococcus epidermidis* is an opportunistic, biofilm forming pathogen that frequently causes hospital-related and medical device-associated infections^15^ and was chosen as the model organism in our studies. Three-dimensional mapping of oxygen gradients within *S. epidermidis* biofilms is important for understanding changes in the spatial heterogeneity of bacterial metabolism and biofilm growth during biofilm development and in response to antimicrobial treatment.

Here we develop a solution-based ratiometric oxygen-sensing platform by coupling the oxygen-sensitive ruthenium complex tris(2,2’-bipyridyl)dichlororuthenium(II) hexahydrate with oxygen-insensitive cadmium selenide zinc sulfide (CdSe/ZnS) quantum dots. This system is integrated with quantitative confocal laser scanning microscopy (CLSM) to spatially map three-dimensional oxygen gradients in *S. epidermidis* biofilms during their development and while undergoing antibiotic treatment. First, we calibrate and validate the oxygen-sensing platform for use within biofilms. Next, we apply the platform to characterize local oxygen concentrations in *S. epidermidis* biofilms during biofilm development and in response to vancomycin treatment. Finally, we evaluate how *S. epidermidis* biofilm structure, metabolic activity, and cellular viability relate to the observed variations in spatial oxygen concentrations.

## 2. Materials and Methods

### 2.1 Solution-based, ratiometric oxygen-sensing platform

The solution-based, ratiometric oxygen-sensing platform consisted of 20 µM tris (2,2’-bipyridyl) dichlororuthenium (II) hexahydrate (Ru(bpy)_3_Cl_2_) (MilliporeSigma, Burlington, MA, USA) and 63 µg/mL water-soluble, Cadmium Selenide/Zinc Sulfide (CdSe/ZnS) quantum dots (Product #HECZW-560, NNCrystal US Corporation, Fayetteville, AR, USA).

Ru(bpy)_3_Cl_2_ is an oxygen sensitive luminophore whose fluorescence is quenched in the presence of oxygen, enabling quantification of dissolved oxygen concentrations of ∼0-8 mg/L in solution^16^. Ru(bpy)_3_Cl_2_ is ∼ 1.2 nm^17^ with an excitation/emission of 452/620 nm^18^, where the fluorescent intensity of Ru(bpy)_3_Cl_2_ decreases linearly with increasing O_2_ concentration in solution^16^. CdSe/ZnS quantum dots are ∼8-9 nm, high-efficiency, water-soluble, fluorescent probes with an excitation/emission of 540/560 nm^19^. The fluorescent intensity of CdSe/ZnS quantum dots has been shown to be stable with varying oxygen concentrations^20^. Coupling oxygen sensitive Ru(bpy)_3_Cl_2_ with the oxygen insensitive CdSe/ZnS quantum dots enabled ratiometric imaging of oxygen concentrations.

### 2.2 Bacterial strains and growth conditions

*S. epidermidis* RP62A was obtained from American Type Culture Collection (ATCC 35984) and is a strong-biofilm former^21^. *S. epidermidis* RP62A was maintained on tryptic soy agar (TSA) or in tryptic soy broth supplemented with 1 wt. % glucose (TSB_g_).

To evaluate the sensitivity of planktonic *S. epidermidis* growth to the two components of the ratiometric oxygen sensor, planktonic *S. epidermidis* RP62A cells were cultured at 37°C and 180 RPM in bacterial culture tubes containing 20 µM Ru(bpy)_3_Cl_2_ and 63 µg/mL water-soluble CdSe/ZnS quantum dots in TSB_g_. OD_600_ measurements of planktonic *S. epidermidis* were taken at 0, 6, 12, 18, and 24 hours to monitor bacterial growth in the presence of the two components of the ratiometric oxygen-sensing platform. Three independent experimental replicates were obtained for each growth condition.

*S. epidermidis* RP62A biofilms were grown by seeding 5x10^5^ cells/mL of an overnight liquid culture into the wells of an 8-well Nunc^TM^ Lab-Tek^TM^ II Chambered Coverglass dish (Thermo Fisher Scientific, Waltham, MA, USA) and culturing the bacteria at 60 RPM and 37°C.

To evaluate the sensitivity of *S. epidermidis* biofilm growth to the two components of the ratiometric oxygen sensor, *S. epidermidis* biofilms were grown at 60 RPM and 37°C with 20 µM Ru(bpy)_3_Cl_2_ and 63 µg/mL water-soluble CdSe/ZnS quantum dots in 96-well plates by seeding individual wells with 5x10^5^ cells/mL in TSB_g_. OD_600_ measurements of biofilms were taken at 0, 6, 12, 18, and 24 hours to assess biofilm growth in the presence of the two components of the ratiometric oxygen-sensing platform. Three independent experimental replicates were obtained for each growth condition.

*S. epidermidis* biofilm development was studied using biofilms grown for 12, 18, and 24 hours. The effect of vancomycin treatment on established *S. epidermidis* biofilms was studied by first growing biofilms for 24 hours, exchanging the growth media for 600 µL TSB_g_ containing 0 µg/mL, 2 µg/mL, 20 µg/mL, or 200 µg/mL vancomycin, and growing the biofilms for an additional 24 hours. These concentrations of vancomycin were chosen as the minimum inhibitory concentration (MIC) of vancomycin for *S. epidermidis* is 2.0 µg/mL^22^, typical trough concentrations of vancomycin for clinical treatment of staphylococcal infections are 10-20 µg/mL^23, 24^, and the minimum biofilm eradication concentration (MBEC) of vancomycin toward *S. epidermidis* biofilms is reported to vary between 4x to greater than 100x more than the MIC, which corresponds to 8 to > 200 µg/mL depending on the strain and growth condition studied^25–27^. Four independent experimental replicates were obtained for each growth condition.

For evaluation of *S. epidermidis* metabolic activity using resuzarin assays, *S. epidermidis* biofilms were grown in 96-well plates by seeding individual wells with 5x10^5^ cells/mL and culturing wells for 12, 18, or 24 hours for the biofilm development study or for 24 hours in TSB_g_, followed by a growth media exchange for 0, 2, 20, or 200 µg/mL vancomycin in TSB_g_ and an additional 24 hours of growth at 60 RPM and 37°C for the vancomycin study. Experiments were performed in biological triplicate, with each growth condition assessed using three technical replicates per experiment (9 total wells per growth condition).

### 2.3 Measurement of dissolved oxygen using oxygen meter

A PreSens Fibox 4 fiber-optic oxygen meter in combination with PSt3 sensor dots (PreSens Precision Sensing GmbH, Regensburg, Germany) was used to measure dissolved oxygen concentrations in calibration solutions and biofilm supernatants. PSt3 sensor dots support the measurement of 0-100% dissolved oxygen or 0-45 mg/L dissolved oxygen with sensitivities of ±0.4% dissolved oxygen at 20.9% O_2_ and ±0.05% dissolved oxygen at 0.2% O_2_. The oxygen meter and PSt3 sensor dots were calibrated with a two-point calibration method using nitrogen-purged deoxygenated water (0% O₂) and air-equilibrated water (21% O₂).

Dissolved oxygen measurements of calibration solutions and biofilm supernatants were conducted in a single well of a 8-well Nunc^TM^ Lab-Tek^TM^ II Chambered Coverglass dish by affixing a PSt3 sensor dot to the side of each well in a chambered coverglass, where the fiber optic probe for measuring dissolved oxygen was positioned at the center of each sensor dot and on the exterior of the plastic side wall of the well during measurement. PSt3 sensor dots were sterilized prior to use by immersion in 70% ethanol for 30 minutes. Sample volumes of 600 µL were used for all measurements in 8-well chambered coverglass dishes to allow for submersion of the PSt3 sensor dots. Affixing PSt3 sensor dots to the wells of the 8-well chambered coverglass dishes did not affect biofilm development as evaluated by OD_600_ measurements of biofilms grown with and without PSt3 sensor dots (Fig. S1).

### 2.4 Calibration of solution-based, ratiometric oxygen-sensing platform for evaluating oxygen concentration in biofilms

The solution-based, ratiometric oxygen-sensing platform was calibrated in solution using a three-point calibration and within alginate hydrogels using a two-point calibration. Fluorescent intensities of the oxygen insensitive CdSe/ZnS quantum dots and oxygen sensitive Ru(bpy)_3_Cl_2_ within each calibration solution or hydrogel were measured using a Leica STELLARIS 8 confocal laser scanning microscope (CLSM) equipped with a 63x, 1.4 NA, oil immersion objective lens. Each component of the calibration solution or hydrogel was imaged using the 63x/1.4 NA objective 5 µm above the coverslip. Four calibration replicates were imaged for each calibration solution or hydrogel. Calibration images were 185 x 185 µm with pixels of 0.361 µm x 0.361 µm.

The linear Stern-Volmer relationship was used to develop oxygen concentration calibration curves using fluorescence intensity ratios between the oxygen sensitive Ru(bpy)_3_Cl_2_ and oxygen insensitive CdSe/ZnS quantum dots, similar to previously established methods for calibration of ratiometric oxygen sensors^28^. The Stern-Volmer relationship describes the quenching of a fluorophore in the presence of the quenching species^29^, in this case oxygen, as follows:

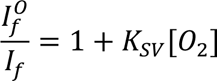

where 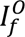 is the fluorescence intensity ratio of Ru(bpy)_3_Cl_2_ to CdSe/ZnS quantum dots under oxygen-free conditions, *I_f_* is the fluorescence intensity ratio of Ru(bpy)_3_Cl_2_ to CdSe/ZnS quantum dots at the concentration of oxygen in solution [*O*_2_], *K_sv_* is the Stern-Volmer constant and [*O*_2_] is the concentration of dissolved oxygen in solution.

An unconfined, solution based, three-point calibration curve was developed using fluorescent intensities from CLSM images of the following calibration solutions: TSB_g_ with dissolved oxygen concentrations adjusted to 0.04 ± 0.02 mg/L, 3.84 ± 0.09 mg/L, and 7.93 ± 0.03 mg/L using sodium sulfite at concentrations of 4 g/L, 0.2 g/L, and 0 g/L, respectively, as sodium sulfite has been previously utilized for removing dissolved oxygen from aqueous solutions^30^. Dissolved oxygen concentrations were verified using the PreSens Fibox 4 fiber-optic oxygen meter. The three-point calibration curve was used to verify the linearity of the Stern-Volner relationship for the chosen probes in the solution-based, ratiometric oxygen-sensing platform.

The two-point calibration curve of the solution-based, ratiometric oxygen-sensing platform in alginate hydrogels was developed to better approximate confinement of the oxygen-sensing probes within a biofilm, similar to the approach used by Jewell et al. for calibrating 163 nm oxygen-sensitive polymeric nanosensors for evaluating oxygen concentration in *Pseudomonas aeruginosa* biofilms^8^. Briefly, alginate hydrogels were formulated by thoroughly mixing a 2 wt% solution of alginic acid sodium salt (Sigma Aldrich, USA) with 63 µg/mL CdSe/ZnS quantum dots and 22 µM Ru(bpy)_3_Cl_2_ and pouring the solution into a 10-mm cylindrical PDMS mold placed on top of agar containing 2% calcium chloride. Next, alginate was allowed to crosslink under vacuum for 30 minutes. Crosslinked alginate hydrogels were then transferred to 8-well chambered coverglass dishes and submerged in 600 µL of oxygen-depleted or oxygen-saturated TSB_g_ prior to CLSM imaging. Oxygen-depleted TSB_g_ (0.013 ± 0.01 mg/L) was achieved by adding 4 g/L sodium sulfite, an oxygen scavenger that reacts with dissolved oxygen to form sodium sulfate and thereby reduce oxygen levels in solution^30^. Oxygen-saturated TSB_g_ (7.92 ± 0.03 mg/L) was established by equilibrating fresh, air saturated TSB_g_ at ambient conditions. Dissolved oxygen concentrations were verified using a Presens Fibox 4 oxygen sensor meter equipped with PSt3 sensor dots. The Stern-Volmer constant from the alginate hydrogel calibration curve was subsequently used to evaluate pixel level oxygen concentrations within biofilms.

### 2.5 Validation of solution-based, ratiometric oxygen-sensing platform for evaluating oxygen concentration in biofilms

The oxygen-sensing platform calibration using alginate hydrogels was validated in biofilms by evaluating dissolved oxygen concentrations in biofilms after modulation of biofilm oxygen content. Oxygen concentrations in *S. epidermidis* biofilms grown for 24-hours were determined using CLSM imaging for untreated biofilms, biofilms with oxygen depleted from their environment chemically by 4 g/L sodium sulfite, or oxygen-rich biofilms that were heat treated at 60°C for 20 minutes followed by media replacement with fresh TSB_g_. Dissolved oxygen concentrations in the biofilm supernatant for untreated biofilms and those with oxygen-depleted and oxygen-rich conditions were independently measured using an oxygen meter prior to imaging.

For each biofilm, dissolved oxygen concentrations were quantified using ratiometric CLSM imaging of an image plane 5 µm above the coverglass. Pixel-level oxygen concentrations were calculated using the established hydrogel calibration curve and the average oxygen concentration within the imaging plane was compared to the dissolved oxygen concentration measured using the oxygen meter.

### 2.6 Confocal laser scanning microscopy (CLSM) of bacterial biofilms

*S. epidermidis* biofilms were labeled with the two components of the solution-based, ratiometric oxygen-sensing platform (20 µM Ru(bpy)_3_Cl_2_, 63 µg/mL CdSe/ZnS quantum dots) to enable mapping of oxygen concentrations within the biofilms as well as 3 µM SYTO^TM^ 9 (Invitrogen, Waltham, MA, USA), a membrane-permeable, fluorescent nucleic acid binding stain, to label bacterial cells within the biofilm. For the vancomycin treatment studies, *S. epidermidis* biofilms were also fluorescently labeled with 3 µM SYTOX^TM^ Blue (Invitrogen, Waltham, MA, USA), a nucleic acid stain that is impermeant to live cells, which enables identification of cells with compromised cell membranes.

*S. epidermidis* biofilms were imaged using a Leica STELLARIS 8 CLSM with a 63x, 1.4 NA, oil immersion objective lens. The oxygen sensitive fluorophore Ru(bpy)_3_Cl_2_ was excited at 440 nm and emission spectra were monitored between 600-640 nm. CdSe/ZnS quantum dots were excited at 540 nm and emission spectra were monitored between 554-574 nm. SYTO^TM^ 9 was excited at 488 nm and emission spectra were monitored between 493-523 nm. SYTOX^TM^ Blue was excited at 405 nm and emission spectra were monitored between 445-480 nm. Five CLSM image volumes of 185 x 185 x 25 µm^3^ with 0.361 µm x 0.361 µm x 1 µm voxels were acquired for each biofilm with four replicates per condition for a total of 20 image volumes per biofilm growth condition. Quantitative image analysis of oxygen concentration and bacterial biomass was performed on the CLSM image volumes.

### 2.7 Quantitative image analysis of biofilm oxygen concentrations

Biofilm dissolved oxygen concentrations were spatially resolved through quantitative image analysis of biofilms labeled with Ru(bpy)_3_Cl_2_ and CdSe/ZnS quantum dots. The alginate hydrogel oxygen concentration calibration curve for the solution-based ratiometric imaging platform was applied to evaluate dissolved oxygen concentrations at individual pixels within the CLSM image volumes. Dissolved oxygen concentration values were calculated and reported for each z-slice within the biofilm up to the maximum biofilm height. The oxygen calibration curve was established for oxygen concentrations between 0 and 8 mg/L. Pixels with fluorescence intensity ratios of Ru(bpy)_3_Cl_2_ to CdSe/ZnS quantum dots that resulted in oxygen concentration values outside of the range of the calibration curve were omitted from analysis (∼3.5% of pixels across all analyzed images). Out-of-range pixels primarily occurred at the extremes of signal intensity and likely resulted from Ru(bpy)_3_Cl_2_ saturation under oxygen-depleted conditions or low-signal-to-noise ratios when fluorophore intensity values of Ru(bpy)_3_Cl_2_ were very low.

Oxygen gradients in the vertical direction (z-direction) were assessed by averaging pixel-level oxygen concentrations at each z-slice of the biofilm image volume, which enabled generation of height-dependent oxygen concentration profiles. The magnitude of the change in oxygen concentration with biofilm height was calculated as Δ[*O*_2_]/Δ*Z*, where the difference between the average oxygen concentration at the top of each biofilm and that at the substrate interface is divided by biofilm height.

### 2.8 Quantitative image analysis of biofilm structure and viability

Biofilm height, biovolume, and porosity were quantified to evaluate biofilm structure using COMSTAT analysis of CLSM image volumes. COMSTAT is a MATLAB-based program for image analysis of biofilm structure^31^. Biofilm height is reported as the average thickness over the entire substratum area of the image volume^31^. Biofilm biovolume^31^ is the total volume of biofilm biomass divided by the biofilm surface area at the substrate surface with units of µm^3^/µm^2^. Biofilm porosity is the ratio of the surface area of biomass void space to the total surface area within an xy image plane. Porosity values were averaged across all z-planes in each image volume up to the maximum height of the biofilm.

For the vancomycin studies, variations in bacterial biomass viability across growth environments were evaluated as the ratio of the dead cell biovolume to total biovolume. The total biofilm biovolume and dead cell biovolume were determined using COMSTAT analysis of the SYTO^TM^ 9 (total bacterial cell biovolume) and SYTOX^TM^ Blue (dead cell biovolume) channels within CLSM image volumes. The total number of pixels with biofilm biomass and dead cells in each image volume were determined using COMSTAT and were multiplied by the size of each voxel (0.361 µm x 0.361 µm x 1 µm) to quantify total biovolume of biofilm biomass or dead cell biovolume. Percentage dead biomass is computed using the ratio of dead cell biovolume to total biovolume.

### 2.9 Evaluation of bacterial biofilm metabolic activity using resazurin assay

Metabolic activity was quantified in biofilms using a resazurin reduction assay. The resazurin reduction assay quantifies cellular respiration and can be used as an indicator of biofilm viability^32^. Following *S. epidermidis* biofilm growth in 96-well plates, 20 µL of 100 µg/mL resazurin in TSB_g_ was added directly to each well of the 96-well plate for a final resazurin concentration of 10 µg/mL. Biofilms were not washed to ensure metabolic activity was captured. The 96-well plates were incubated at 37°C for 20 minutes to allow the resazurin to be reduced to resorufin. The fluorescence intensity of the wells was captured using a VICTOR Nivo^TM^ multimode plate reader (PerkinElmer Inc., Waltham, MA, USA), where the resorufin was excited with a 540/30 nm filter and emission was captured with a 580/20 nm filter. Resazurin assay fluorescent intensity values were normalized to 12-hour samples for the biofilm development studies and to biofilms grown without vancomycin treatment (0 µg/mL) for the vancomycin treatment studies to assess the relative metabolic activity of biofilms.

### 2.10 Statistical analysis

Statistical analyses were performed using GraphPad Prism (version 10.6.1, GraphPad Software, Boston, MA, USA). Data are reported as mean ± standard error of the mean unless otherwise noted. Comparisons between dissolved oxygen concentrations measured using the solution-based oxygen-sensing platform and the bulk oxygen meter were analyzed using a two-way ANOVA followed by Tukey’s multiple comparisons test. Pair-wise comparisons of *S. epidermidis* planktonic and biofilm growth in the absence and presence of Ru(bpy)_3_Cl_2_ and CdSe/ZnS quantum dots at each growth time (0, 6, 12, 18, and 24 hours) were analyzed using two-way ANOVA with Bonferroni’s multiple comparisons post hoc test. For datasets with unequal variance (dissolved oxygen concentrations in biofilm control conditions (Fig. 2), rates of change in oxygen concentration with biofilm height (Δ[O₂]/Δz), bacterial metabolic activity, biofilm thickness, biovolume, porosity, and dead biovolume), statistical significance was determined using Brown-Forsythe and Welch ANOVA with Dunnett’s T3 multiple comparisons test. For all statistical analyses, significance was defined as: *ns* (not significant), \**P* ≤ 0.05, \*\**P* ≤ 0.01, \*\*\**P* ≤ 0.001, and \*\*\*\**P* ≤ 0.0001.

**Figure 1.**
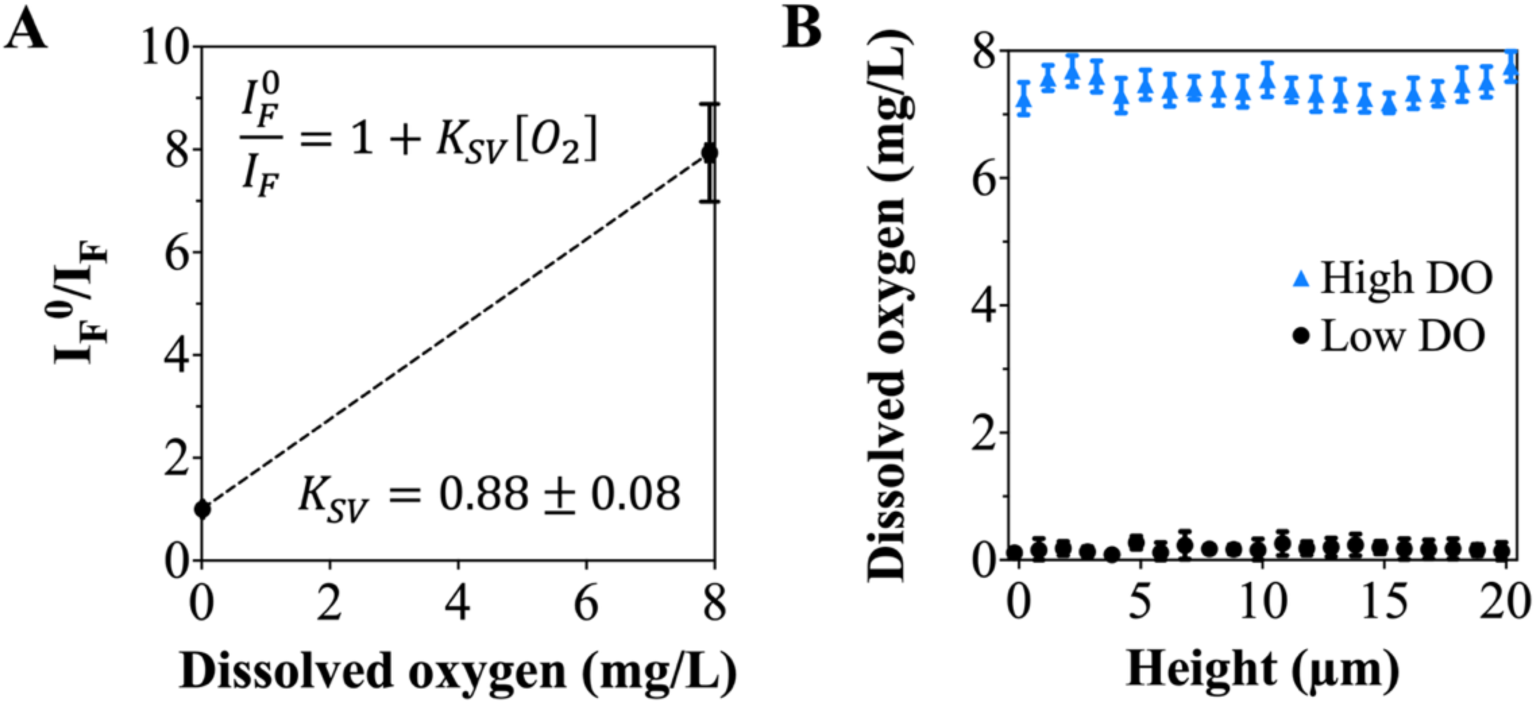
Calibration of oxygen-sensing platform in alginate hydrogels. **(A)** Ratiometric dissolved oxygen calibration curve for oxygen-sensing platform in alginate hydrogels with fluorescence intensity ratio (I_F_^0^/I_F_) plotted against dissolved oxygen concentration (mg/L). **(B)** Average dissolved oxygen concentrations in xy-image planes from 0-20 µm within alginate hydrogels in solutions with low and high dissolved oxygen conditions, where low dissolved oxygen concentrations are achieved using 4 g/L sodium sulfite.

**Figure 2.**
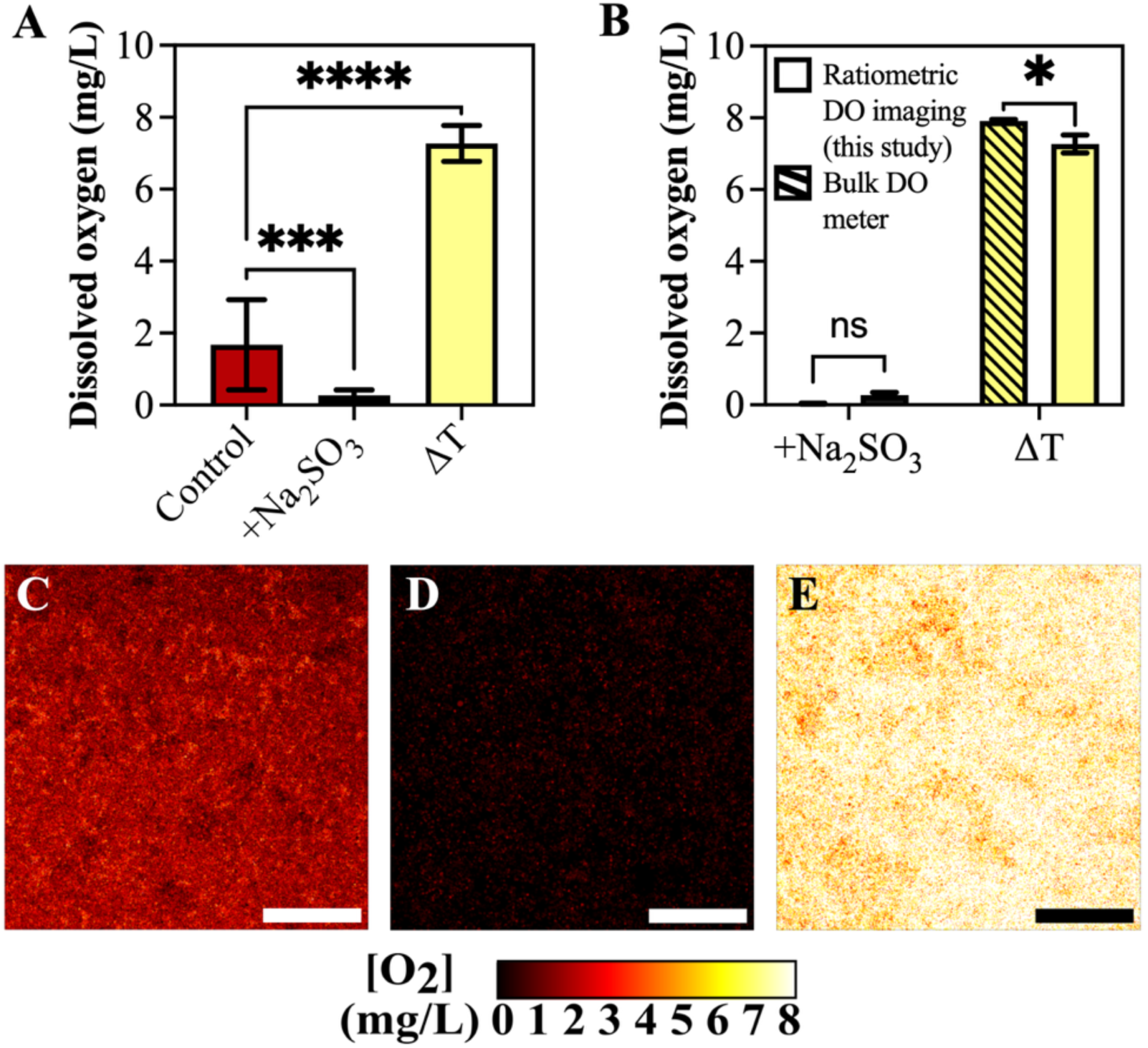
Validation of oxygen-sensing platform calibrated in alginate hydrogels for evaluating dissolved oxygen concentrations in *S. epidermidis* biofilms. **(A)** Quantification of average dissolved oxygen concentration at 5 µm above substrate interface within *S. epidermidis* RP62A biofilms grown for 24 h at 37 °C (control), *S. epidermidis* RP62A biofilms with low dissolved oxygen concentrations achieved by the addition of 4 g/L sodium sulfite to biofilms after 24 h (+Na_2_SO_3_), and *S. epidermidis* RP62A biofilms with high dissolved oxygen concentrations achieved by bacterial thermal inactivation at 60°C followed by TSB_g_ media replacement (ΔT). **(B)** Dissolved oxygen concentrations measured at 5 µm above the substrate with the solution-based, ratiometric imaging platform and in the biofilm supernatant with the bulk dissolved oxygen meter for *S. epidermidis* RP62A biofilms with low (+Na_2_SO_3_) and high dissolved oxygen concentrations (ΔT). (**C-E**) Pseudo-color oxygen concentration maps of *S. epidermidis* RP62A biofilms **(C)** grown for 24-hours (control), **(D)** with low dissolved oxygen concentrations (+Na_2_SO_3_), and **(E)** with high dissolved oxygen concentrations (ΔT). All dissolved oxygen maps and dissolved oxygen concentrations reported are from 5 µm above substrate interface for biofilm growth. Scale bars: 50 µm. Statistical significance for **(A)** was determined by a Brown-Forsythe and Welch ANOVA with Dunnett’s T3 multiple comparisons test and for **(B)** by a two-way ANOVA with Tukey’s multiple comparisons test (*ns*: not significant, \**P* < 0.05, \*\**P* < 0.01, \*\*\**P* < 0.001, \*\*\*\**P* < 0.0001).

## 3. Results

### 3.1 Calibration and validation of solution-based, ratiometric oxygen sensing platform for detecting dissolved oxygen in *Staphylococcus epidermidis* biofilms

The solution-based, ratiometric oxygen-sensing platform consisting of oxygen-sensitive, Ru(bpy)_3_Cl_2_ and oxygen-insensitive, water-soluble CdSe/ZnS quantum dots is calibrated over a dissolved oxygen range of 0-8 mg/L in alginate hydrogels and in solution using the linear Stern-Volmer relationship. The fluorescence intensity of Ru(bpy)_3_Cl_2_ decreases linearly as dissolved oxygen concentration increases (Fig. 1A, Supp. Fig. S2). The Stern-Volmer constant, *K_SV_*, is 0.88 ± 0.08 L/mg for the calibration performed in the alginate hydrogel and 0.31 ± 0.02 L/mg for the calibration performed in solution, where the uncertainty represents the standard error of the fitted slope. The alginate hydrogel calibration curve is used for determining dissolved oxygen concentration in biofilms due to the similarities between the confined structures of the hydrogels and biofilms. Both high and low dissolved oxygen concentrations are stable and spatially uniform within alginate hydrogels in oxygen-depleted and oxygen-saturated media throughout the first 20 µm of the hydrogel, which validates probe performance throughout the hydrogel height within a defined three-dimensional matrix (Fig. 1B).

Dissolved oxygen concentrations were modulated within *S. epidermidis* biofilms and measured using the oxygen-sensing platform calibrated using alginate hydrogels. *S. epidermidis* biofilms with oxygen depleted from the environment by sodium sulfite contain average dissolved oxygen concentrations of 0.27±0.07 mg/L at 5 µm above the coverslip (Fig. 2A). *S. epidermidis* biofilms with oxygen-rich environments achieved by first thermally inactivating cells within the biofilms and then replacing media with fresh bacterial growth media contained average dissolved oxygen concentrations of 7.27 ± 0.26 mg/L at 5 µm above the coverslip (Fig. 2A). *S. epidermidis* biofilms without modulated dissolved oxygen concentrations contain average dissolved oxygen concentrations of 1.68 ± 0.28 mg/L at 5 µm above the coverslip (Fig. 2A). Thus, modulation of dissolved oxygen within *S. epidermidis* biofilms was successfully achieved. Oxygen meter measurements of dissolved oxygen concentration in the supernatant of oxygen-rich and oxygen-depleted biofilms are 7.92±0.04 mg/L and 0.030±0.01 mg/L, respectively, representing an 8-10% difference between the imaging-based measurements and oxygen meter measurements of oxygen concentration within biofilms with modulated oxygen content (Fig. 2B). Pseudo-color oxygen maps of control biofilms, biofilms with low dissolved oxygen (0 mg/mL) and biofilms with high dissolved oxygen (7.9 mg/mL) at 5 µm above the substrate interface show that the modulated oxygen environments result in relatively uniform oxygen concentrations as compared to the control biofilms (Fig. 2C-E). Thus, physiologically relevant dissolved oxygen concentrations are reliably measured in complex *S. epidermidis* biofilm structures using our oxygen-sensing platform calibrated within alginate hydrogels.

*S. epidermidis* planktonic and biofilm growth are not significantly altered by ratiometric oxygen-sensing platform components after 24 hours. OD_600_ measurements of planktonic *S. epidermidis* RP62A grown in the presence of the two oxygen-sensing components for 24 hours showed no significant differences to untreated controls (Fig. 3A). Similarly, *S. epidermidis* RP62A biofilm biomass after 24 hours was not significantly altered by growth with the two oxygen-sensing components (Fig. 3B), indicating minimal effects of the oxygen-sensing platform on both planktonic and biofilm growth of *S. epidermidis* RP62A.

**Figure 3.**
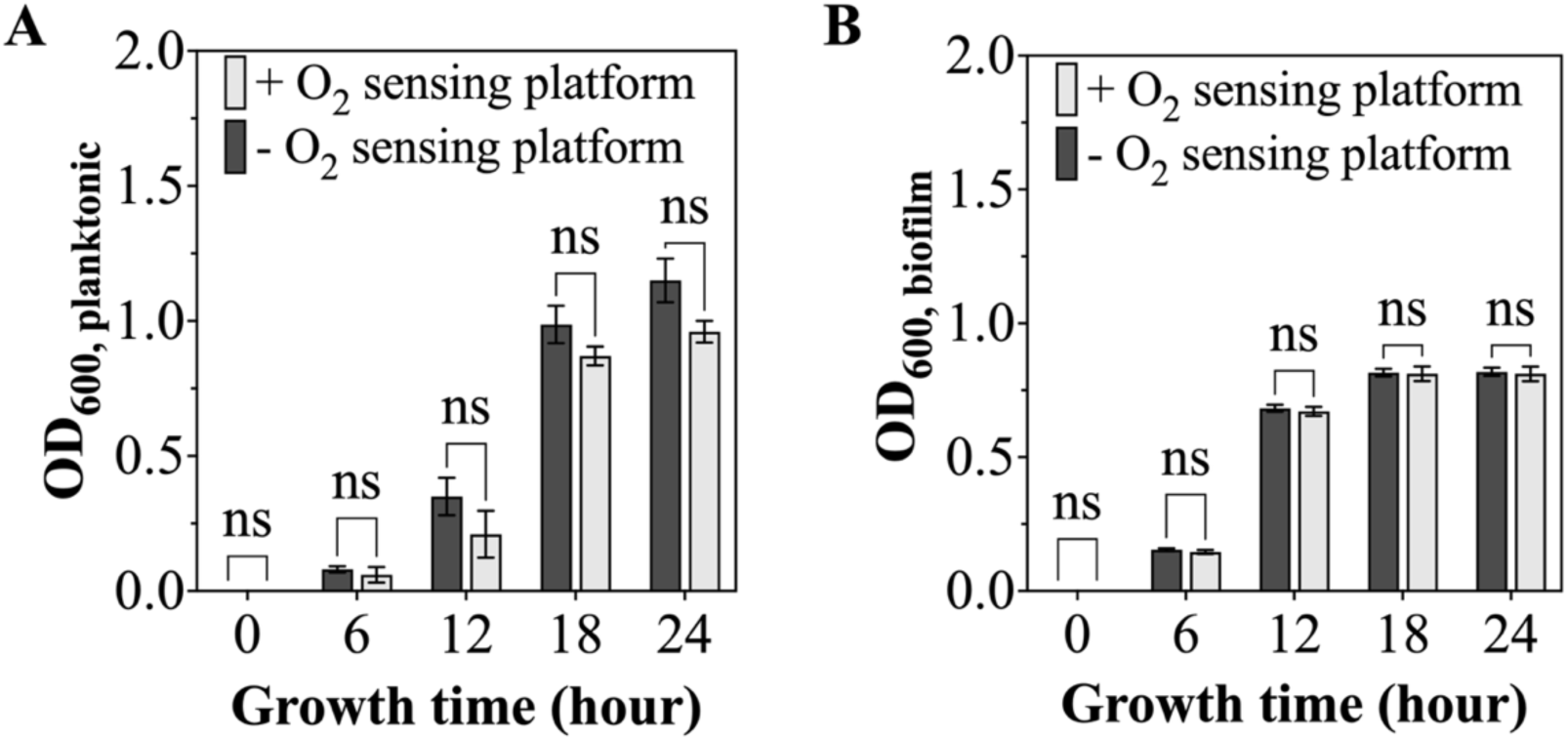
Effect of oxygen-sensing platform components on *S. epidermidis* planktonic and biofilm growth. Optical density measurements of *S. epidermidis* RP62A **(A)** planktonic cells and **(B)** biofilms after at 0, 6, 12, 18 and 24 hours of growth without (- O_2_ sensing platform) or with (+ O_2_ sensing platform) Ru(bpy)_3_Cl_2_ and CdSe/ZnS quantum dots. Statistical significance was determined by a two-way ANOVA with Bonferroni’s multiple comparisons test (*ns*: not significant).

### 3.2 Oxygen gradient formation during *S. epidermidis* biofilm development

Three-dimensional dissolved oxygen maps of *S. epidermidis* biofilm development show that dissolved oxygen concentrations decrease throughout the biofilm structure from 12 to 24 hours of biofilm growth (Fig. 4A-C), where the average dissolved oxygen concentration within the biofilm decreases from 2.62±0.22 mg/L after 12 hours to 2.26±0.13 mg/L after 18 hours to 2.02±0.47 mg/L after 24 hours (Fig. 4D). Dissolved oxygen concentrations determined using sub-micron scale imaging of biofilms differ most significantly from dissolved oxygen concentrations measured in the biofilm supernatant with an oxygen meter at 12 hours of biofilm development, where bulk dissolved oxygen concentrations are 3.50±0.23 mg/L at 12 hours, 2.28±0.14 mg/L at 18 hours, and 2.04±0.21 mg/L at 24 hours (Fig. 4D).

**Figure 4.**
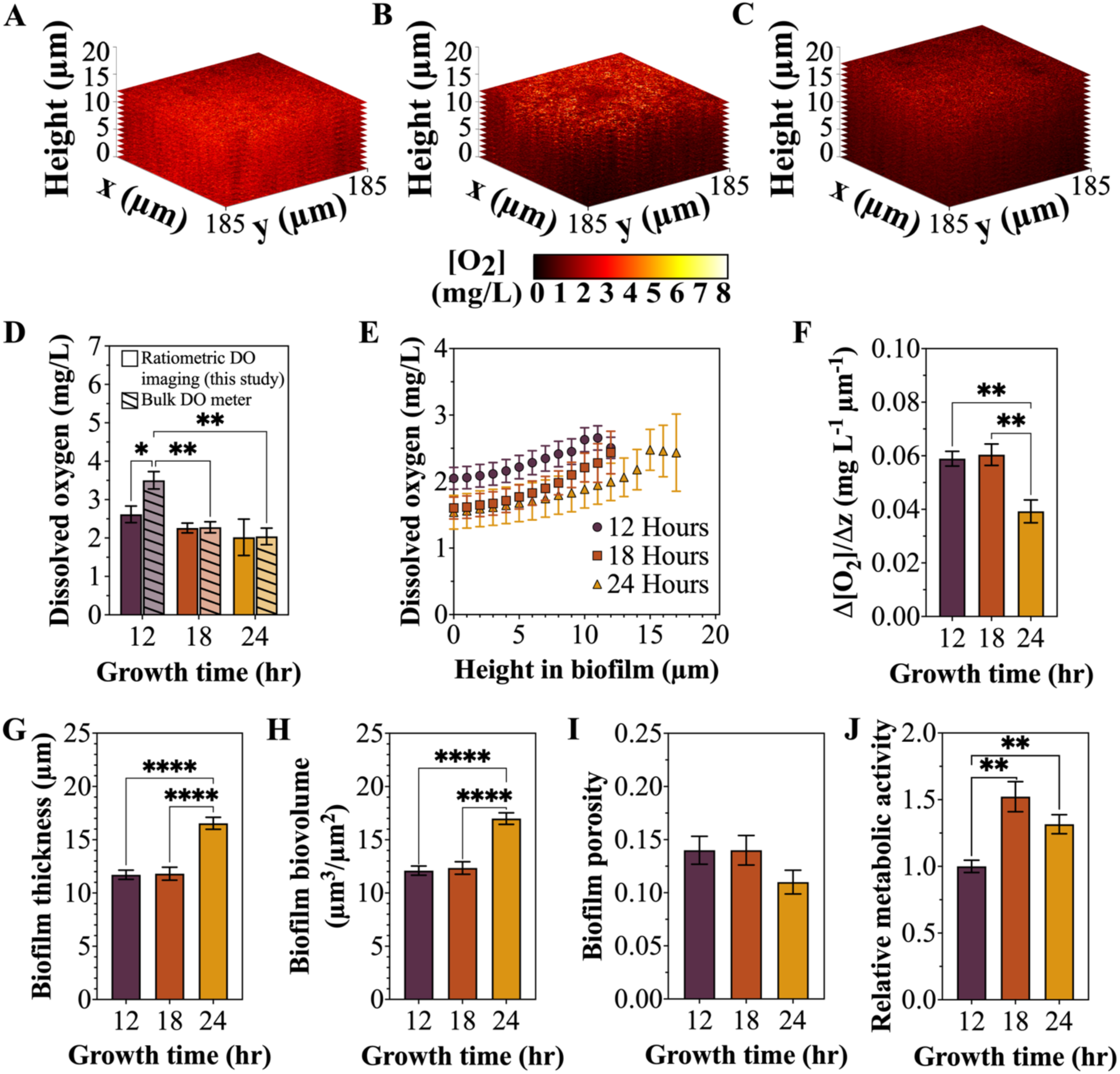
Spatiotemporal changes in dissolved oxygen concentration, biofilm structure, and bacterial metabolic activity during *S. epidermidis* biofilm development. **(A-C)** Three-dimensional *S. epidermidis* biofilm oxygen concentration maps from ratiometric imaging using oxygen-sensing platform coupled with CLSM after **(A)** 12-hours, **(B)** 18-hours, and **(C)** 24 hours of biofilm growth. **(D)** Average dissolved oxygen concentrations measured with the solution-based, ratiometric imaging platform and the bulk dissolved oxygen meter after 12, 18, and 24 hours of *S. epidermidis* RP62A biofilm growth. **(E)** Dissolved oxygen concentration throughout biofilm height after 12, 18, and 24 hours of biofilm growth. **(F)** Rate of change in oxygen concentration with biofilm height (Δ[O₂]/Δz) after 12-, 18-, and 24-hours of biofilms growth. (G-I) *S. epidermidis* biofilm **(G)** thickness, **(H)** biovolume, and **(I)** porosity from 12 to 24 hours of biofilm growth. **(J)** Bacterial metabolic activity relative to 12-hours *of S.* epidermidis biofilm growth measured using a resazurin assay after 12, 18, and 24-hours of biofilm growth. Statistical significance for **(D)** was determined by a two-way ANOVA with Tukey’s multiple comparisons test and **(F-J)** were determined by a Brown-Forsythe and Welch ANOVA with Dunnett’s T3 multiple comparisons test (*ns*: not significant, \**P* < 0.05, \*\**P* < 0.01, \*\*\**P* < 0.001, \*\*\*\**P* < 0.0001).

Dissolved oxygen concentrations also vary with height within the biofilm, where *S. epidermidis* RP62A biofilms consistently have the lowest dissolved oxygen concentrations at the substrate interface where initial bacterial adhesion occurs (Fig. 4A-C, 4E). Over 24-hours of biofilm development, oxygen availability decreases locally at each height within the biofilm (Fig. 4A-C, 4E). At the *S. epidermidis* biofilm substrate interface, dissolved oxygen decreases from 2.05±0.16 mg/L at 12 hours to 1.60±0.17 mg/L at 18 hours to 1.54±0.25 mg/L at 24 hours, representing a 0.51 mg/L decrease in average dissolved oxygen at the growth substrate from 12 to 24 hours of biofilm development (Fig. 4E). *S. epidermidis* biofilm thickness after 12 hours is 11.7 ± 1.9 µm; at 12 µm above the substrate interface, the dissolved oxygen concentration decreases from 2.51±0.16 mg/L at 12 hours to 2.44±0.32 mg/L at 18 hours to 2.00±0.28 mg/L at 24 hours, representing a 0.51 mg/L decrease in average dissolved oxygen from 12 to 24 hours at 12 µm above the substrate interface (Fig. 4E). After 24 hours, *S. epidermidis* biofilm thickness is 16.5 ± 2.5 µm and the average dissolved oxygen concentration 17 µm above the substrate interface at 24 hours is 2.44±0.58 mg/L (Fig. 4E). Thus, the average dissolved oxygen concentration at the top of the 24-hour *S. epidermidis* RP62A biofilms is 0.09 mg/mL lower than the dissolved oxygen concentration at the top of *S. epidermidis* biofilms grown for 12-hours. Thus, changes in dissolved oxygen concentration vary with height above the substrate interface throughout biofilm development. Dissolved oxygen concentrations in the z-direction vary with biofilm growth time, where concentration gradients are similar and larger after 12-hour and 18-hour growth (0.059±0.002 mg/L-µm and 0.060±0.004 mg/L-µm, respectively), as compared to after 24-hours of growth (0.039±0.004 mg/L-µm), representing a 33-35% decrease in dissolved oxygen concentration change in the z-direction as compared to 12-hour and 18-hour biofilms (Fig. 4F).

As average dissolved oxygen concentrations decrease within *S. epidermidis* biofilms during biofilm development from 12 to 24 hours, biofilm thickness and biovolume increase and bacterial metabolic activity fluctuates. Biofilm height is 11.7±1.9 µm after 12 hours, 11.8±2.7 µm after 18 hours, and 16.5±2.5 µm after 24 hours of biofilm development (Fig. 4G). Increases in biofilm height are coupled with increases in biofilm biomass accumulation, where biovolume is 12.1±1.9 µm^3^/µm^2^ after 12 hours, 12.4±2.6 µm^3^/µm^2^ after 18 hours, and 17.0±2.4 µm^3^/µm^2^ after 24 hours (Fig. 4H). Average biofilm porosity does not change significantly from 12 to 24 hours of *S. epidermidis* RP62A biofilm growth (Fig. 4I). Dissolved oxygen concentration and structural changes in *S. epidermidis* RP62A biofilms from 12-24 hours of growth are accompanied by increases in bacterial metabolic activity, where resazurin assays indicate that relative metabolic activity is 1.5 times higher after 18 hours, and 1.3 times higher after 24 hours of biofilm growth as compared to after 12-hours of biofilm growth (Fig. 4J).

### 3.3 Oxygen gradients in *S. epidermidis* biofilms after vancomycin treatment

Three-dimensional dissolved oxygen maps of *S. epidermidis* biofilms after treatment with 0-200 µg/mL vancomycin show that dissolved oxygen concentrations increase with increases in vancomycin concentration (Fig. 5A-D), where average dissolved oxygen concentration increases from 1.84±0.29 mg/L with 2 µg/mL vancomycin to 3.10±0.19 mg/L with 20 µg/mL vancomycin, and to 3.43±0.48 mg/L with 200 µg/mL vancomycin treatment (Fig. 5E). The dissolved oxygen concentrations determined using sub-micron scale imaging of biofilms are consistently lower than dissolved oxygen concentrations measured in the biofilm supernatant with an oxygen meter after vancomycin treatment with statistically significant differences between the measurements for biofilms treated with 20 and 200 µg/mL vancomycin, where bulk dissolved oxygen concentrations are 2.45±0.26 mg/L for 2 µg/mL vancomycin, 4.26±0.10 mg/L for 20 µg/mL vancomycin, and 6.35±0.33 mg/L for 200 µg/mL vancomycin (Fig. 5E).

**Figure 5.**
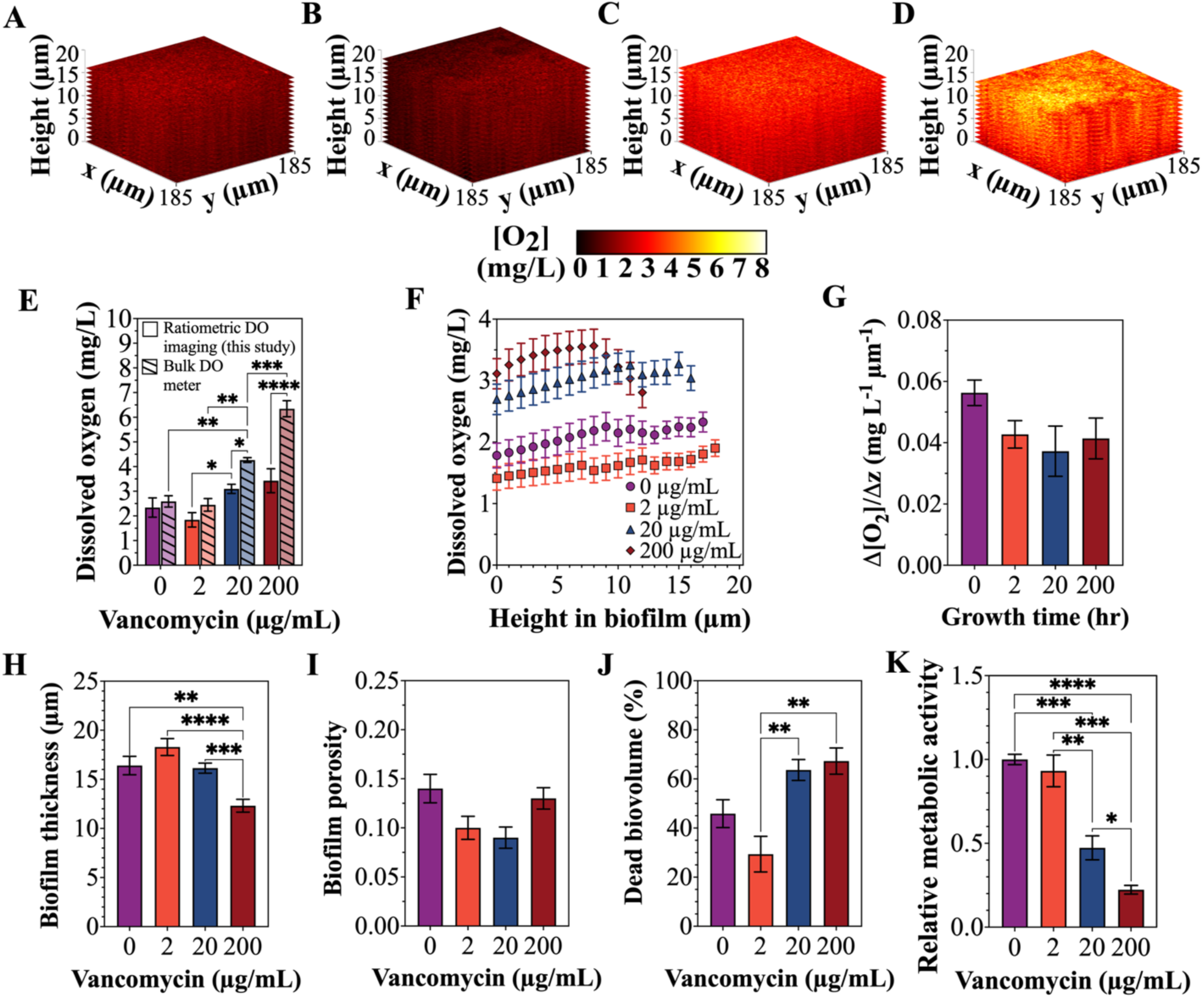
Effect of vancomycin treatments on microscale dissolved oxygen concentrations, biofilm structure, and bacterial metabolic activity of *S. epidermidis* biofilms. **(A-D)** Three-dimensional *S. epidermidis* RP62A biofilm oxygen concentration maps from ratiometric imaging using oxygen-sensing platform coupled with CLSM after treatment with **(A)** 0 µg/mL, **(B) 2** µg/mL, **(C)** 20 µg/mL, and **(D)** 200 µg/mL vancomycin. **(E)** Average dissolved oxygen concentrations measured with the solution-based, ratiometric imaging platform and the bulk dissolved oxygen meter after treatment of *S. epidermidis* RP62A biofilms with 0-200 µg/mL vancomycin. **(F)** Dissolved oxygen concentration throughout biofilm height after treatment with 0-200 µg/mL vancomycin. **(G)** Rate of change in oxygen concentration with biofilm height (Δ[O₂]/Δz) after treatment with 0-200 µg/mL vancomycin. (H-J) *S. epidermidis* biofilm **(H)** thickness, **(I)** porosity, and **(J)** percentage dead biovolume after treatment with 0-200 µg/mL vancomycin. **(K)** Bacterial metabolic activity relative to untreated *S.* epidermidis biofilms measured using a resazurin assay after treatment with 0-200 µg/mL vancomycin. Statistical significance for **(E)** was determined by a two-way ANOVA with Tukey’s multiple comparisons test and **(G-K)** were determined by a Brown-Forsythe and Welch ANOVA with Dunnett’s T3 multiple comparisons test (ns- not significant, *p < 0.05, **p < 0.01, ***p < 0.001, ****p < 0.0001).

Dissolved oxygen concentrations after vancomycin treatment vary with height within the biofilms in a dose-dependent manner, where *S. epidermidis* RP62A biofilms treated with 2 µg/mL vancomycin have lower oxygen concentrations at all spatial positions within the biofilm, and biofilms treated with 20 µg/mL or 200 µg/mL vancomycin have increased oxygen concentrations throughout the biofilm relative to biofilms without antibiotic treatment (Fig. 5A-D, 5F). At the *S. epidermidis* biofilm substrate interface, dissolved oxygen concentration is 1.78±0.20 mg/L with 0 µg/mL vancomycin, 1.41±0.19 mg/L with 2 µg/mL vancomycin, 2.70±0.25 mg/L with 20 µg/mL vancomycin, and 3.11±0.25 mg/L with 200 µg/mL vancomycin treatment (Fig. 5F), corresponding to a 21% decrease, 52% increase and 75% increase in average dissolved oxygen concentration as compared to untreated biofilms after treatment with 2, 20, and 200 µg/mL vancomycin, respectively. The change of biofilm dissolved oxygen concentration with height does not vary significantly after vancomycin treatment, where average oxygen concentration gradients in the z-direction are 0.043±0.004 mg/L-µm , 0.037±0.008 mg/L-µm, and 0.041±0.007 mg/L-µm after treatment with 2, 20, and 200 µg/mL vancomycin, respectively (Fig. 5G), which corresponds to a 23-27% reduction in the dissolved oxygen concentration gradient within *S. epidermidis* biofilms after vancomycin treatment relative to untreated biofilms.

Variations in *S. epidermidis* biofilm dissolved oxygen concentrations with vancomycin dosage correspond to changes in biofilm thickness, viability and metabolic activity. *S. epidermidis* biofilm height is similar for biofilms treated with 0-20 µg/mL vancomycin and significantly decreases after treatment with 200 µg/mL vancomycin (Fig. 5H), while biofilm porosity does not vary significantly between vancomycin treatments (Fig. 5I). The proportion of dead cell biomass increases most significantly between vancomycin treatment dosages of 2 µg/mL and 200 µg/mL, where dead cell biomass increases from 29 to 67% (Fig. 5J). *S. epidermidis* biofilms with 20 µg/mL and 200 µg/mL vancomycin treatments have the highest fractions of dead cell biomass (64 and 67%, respectively, Fig. 5J), which corresponds with the highest local dissolved oxygen concentrations after vancomycin treatment (Fig. 5A-D, 5F). While all *S. epidermidis* biofilms maintain metabolic activity after vancomycin treatments of 2-200 ug/mL, bacterial metabolic activity is 53% and 78% lower than in untreated biofilms after treatment with 20 µg/mL and 200 µg/mL vancomycin, respectively (Fig. 5K).

## 4. Discussion

This research establishes a solution-based ratiometric imaging platform for fluorescently mapping dissolved oxygen concentrations in *S. epidermidis* biofilms. The imaging platform consists of two nanoscale components, Ru(bpy)_3_Cl_2_ and CdSe/ZnS quantum dots, calibrated in alginate hydrogels to mimic probe confinement within biofilms (Fig. 1) and validated in biofilms with modulated oxygen concentrations (Fig. 2). The fluorescence intensity and quenching behavior of the ratiometric probes can be altered by the surrounding medium due to changes in light scattering, absorption, and probe-media interactions^33, 34^; thus, performing calibration within a hydrogel instead of in solution improves the accuracy of oxygen quantification by accounting for shifts in scattering due to matrix confinement. In addition, the two components of the ratiometric imaging platform are found to not effect viability of *S. epidermidis* planktonic cells or biofilms (Fig. 3), which is significant for probe use with living systems and suggests the probes do not affect bacterial and biofilm growth. Finally, coupling the solution-based imaging platform with quantitative CLSM enables sub-micron resolution, three-dimensional oxygen mapping of biofilms during biofilm development (Fig. 4) and after antibiotic treatment (Fig. 5).

The small size of Ru(bpy)_3_Cl_2_ (∼1.2 nm) and CdSe/ZnS quantum dots (8-9 nm) overcomes key limitations of traditional oxygen micro- and nano-sensor technologies that require either incorporation of larger oxygen-sensing probes (200-1000 nm^7-9^) during the initial stages of biofilm development^6^, two-dimensional planar optodes that lack three-dimensional spatial resolution^6^, or bulk electrodes that perturb biofilm structure^6^. In addition, while ratiometric solution-based sensors have been used previously to map other physicochemical gradients in biofilms, including pH^35–38^ and heavy metals^39^, high-spatial resolution oxygen gradient mapping in biofilms was limited prior to this study. The simplicity of adding these two probes directly to biofilms for non-perturbative *in situ* measurements of biofilms is an additional advantage of our platform.

Using the solution-based platform, *S. epidermidis* biofilm dissolved oxygen concentrations are mapped with sub-micron resolution during biofilm development from 12-24 hours. Local oxygen concentrations were depleted over time, reaching an average concentration of 2.02±0.47 mg/L after 24 hours of biofilm growth, with the lowest dissolved oxygen concentrations consistently observed closest to the substrate interface. Resolving the most anoxic conditions at the substrate interface was also found in *S. aureus* biofilms with dissolved oxygen profiles mapped with height using 1000 nm probes for sensing oxygen^7^. However, our results differ from a prior study of oxygen concentration changes during *S. epidermidis* biofilm development where microelectrode measurements of the supernatant of biofilms grown in 96-well plates showed dissolved oxygen concentrations that reduce to nearly 0 mg/mL oxygen after just 6 hours of growth^40^ as low concentrations of dissolved oxygen remain present throughout 24-hours of biofilm development in our study likely due to differences in biofilm growth techniques between the two studies.

Quantification of *S. epidermidis* biofilm dissolved oxygen concentration changes in response to antibiotic treatment is important for evaluating the ratiometric imaging platform as a method for understanding spatial variations in cellular metabolism within biofilms after antibiotic treatment. Quantification of biofilm dissolved oxygen concentration changes in response to antibiotic treatment can help reveal bacterial survivability in response to treatment as well as local regions of the biofilm that are most valuable to target with additional antimicrobial therapeutics. Here we find that while oxygen concentration within biofilms increased in a dose-dependent manner with increasing vancomycin treatment concentrations, height-dependent variations in oxygen gradients and overall cellular metabolic activity remained across all vancomycin concentrations studied (2, 20, 200 µg/mL). In a study of *P. aeruginosa* biofilms using 163 nm oxygen probes, antibiotic-induced metabolic changes were mapped in a similar manner with lower spatial resolution^8^.

In both the study of oxygen gradients during *S. epidermidis* biofilm development and after vancomycin treatment, there were variations in the differences between the average imaging-based dissolved oxygen concentrations and the dissolved oxygen concentration measurements of the biofilm supernatant using the bulk oxygen meter. This is likely due to the large volume of media above the biofilm and the accumulation of any metabolic byproducts that have diffused out of the biofilm being accounted for in the bulk measurement, while dissolved oxygen concentration is only reported up to the maximum height of the biofilm when performing quantitative oxygen mapping using the ratiometric imaging platform. This highlights the importance of considering sub-micron three-dimensional spatial variations in dissolved oxygen profiles within biofilms for resolving changes in dissolved oxygen, a common proxy for biofilm metabolic activity.

Our validated and tested solution-based ratiometric imaging platform can be applied to future studies to advance the understanding of the emergence of oxygen gradients in additional biofilm growth environments, biofilms formed by other bacterial species, or within multispecies biofilms. The platform is also promising for evaluating antibiofilm efficacy of antibiotics with efficacies that vary with the metabolic state of bacteria, such as aminoglycosides^41^, as stages of biofilm development with higher oxygen availability throughout biofilm structures could be targeted with the oxygen-dependent antibiotics. In addition, the platform can be applied for evaluating how emerging antibiofilm therapeutics (e.g. phage^42^, antimicrobial peptides^43^) modulate the spatial distribution of oxygen availability within biofilms as well as for monitoring biofilm survivability and recovery after the removal of a therapeutic agent.

## 5. Conclusions

This work developed a non-perturbative, solution-based, ratiometric imaging platform that enables *in situ,* sub-micron, quantitative mapping of oxygen gradients within *S. epidermidis* biofilms. The oxygen-sensing platform composed of oxygen sensitive Ru(bpy)_3_Cl_2_ and oxygen insensitive CdSe/ZnS quantum dots is calibrated in alginate hydrogels to enable robust quantification of oxygen gradients within biofilms. We find that *S. epidermidis* biofilm dissolved oxygen concentrations decrease significantly during biofilm development alongside increases in bacterial biomass and bacterial metabolic activity with the most anoxic regions consistently at the substrate interface. In addition, there is a dose-dependent increase in dissolved oxygen concentrations within *S. epidermidis* biofilms after vancomycin treatment (2-200 µg/mL) that corresponds with increased dead cell biovolume and reductions in biofilm metabolic activity. Our nanoscale solution-based oxygen sensing platform can be applied to quantify and assess the interplay between biofilm oxygen gradients, structure, and metabolic activity across other bacterial strains and growth environments or to evaluate the efficacy of novel antimicrobial strategies.

## Author Contributions

This manuscript was written through contributions of all authors. P. J. Michalik performed: Writing – original draft, Writing – review & editing, Visualization, Validation, Software, Methodology, Investigation, Formal analysis, Data curation, and Conceptualization. E. J. Stewart performed: Writing – original draft, Writing – review & editing, Visualization, Validation, Supervision, Project administration, Methodology, Funding acquisition, Formal analysis, and Conceptualization. All authors have given approval for the final version of the manuscript.

## Supporting Information

The supporting information file includes additional results for validating experimental methods including the effect of PSt3 sensor dots on biofilm growth and a solution-based calibration for the oxygen sensing platform.

## Acknowledgements

This work was supported by startup funds from Worcester Polytechnic Institute.

## References

(1) Hall-Stoodley, L.; Costerton, J. W.; Stoodley, P. Bacterial biofilms: from the natural environment to infectious diseases. Nat Rev Microbiol 2004, 2 (2), 95–108. DOI: 10.1038/nrmicro821.

(2) Stewart, P. S.; Franklin, M. J. Physiological heterogeneity in biofilms. Nature Reviews Microbiology 2008, 6 (3), 199–210. DOI: 10.1038/nrmicro1838.

(3) Jo, J.; Price-Whelan, A.; Dietrich, L. E. P. Gradients and consequences of heterogeneity in biofilms. Nat Rev Microbiol 2022, 20 (10), 593–607. DOI: 10.1038/s41579-022-00692-2.

(4) Bridier, A.; Briandet, R. Microbial Biofilms: Structural Plasticity and Emerging Properties. Microorganisms 2022, 10 (1). DOI: 10.3390/microorganisms10010138.

(5) Crabbé, A.; Jensen, P. Ø.; Bjarnsholt, T.; Coenye, T. Antimicrobial Tolerance and Metabolic Adaptations in Microbial Biofilms. Trends in Microbiology 2019, 27 (10), 850–863. DOI: 10.1016/j.tim.2019.05.003.

(6) Saccomano, S. C.; Jewell, M. P.; Cash, K. J. A review of chemosensors and biosensors for monitoring biofilm dynamics. Sensors and Actuators Reports 2021, 3, 100043. DOI: 10.1016/j.snr.2021.100043.

(7) Acosta, M. A.; Velasquez, M.; Williams, K.; Ross, J. M.; Leach, J. B. Fluorescent silica particles for monitoring oxygen levels in three-dimensional heterogeneous cellular structures. Biotechnology and Bioengineering 2012, 109 (10), 2663–2670. DOI: 10.1002/bit.24530.

(8) Jewell M, P.; Galyean A. A.; Harris, J.K.; Zemanick, E. T.; Cash, K. J. Luminescent Nanosensors for Ratiometric Monitoring of Three-Dimensional Oxygen Gradients in Laboratory and Clinical *Pseudomonas aeruginosa* Biofilms. Applied and Environmental Microbiology 2019, 85 (20), e01116–01119. DOI: 10.1128/AEM.01116-19).

(9) Huynh, G. T.; Tunny, S. S.; Frith, J. E.; Meagher, L.; Corrie, S. R. Organosilica Nanosensors for Monitoring Spatiotemporal Changes in Oxygen Levels in Bacterial Cultures. ACS Sensors 2024, 9 (5), 2383–2394. DOI: 10.1021/acssensors.3c02747.

(10) Yoshihara, T.; Hirakawa, Y.; Hosaka, M.; Nangaku, M.; Tobita, S. Oxygen imaging of living cells and tissues using luminescent molecular probes. Journal of Photochemistry and Photobiology C: Photochemistry Reviews 2017, 30, 71–95. DOI: 10.1016/j.jphotochemrev.2017.01.001.

(11) Wang, X.-d.; Wolfbeis, O. S. Optical methods for sensing and imaging oxygen: materials, spectroscopies and applications. Chemical Society Reviews 2014, 43 (10), 3666–3761, 10.1039/C4CS00039K. DOI: 10.1039/C4CS00039K.

(12) Kühl, M.; Rickelt, L. F.; Thar, R. Combined Imaging of Bacteria and Oxygen in Biofilms. Applied and Environmental Microbiology 2007, 73 (19), 6289–6295. DOI: doi:10.1128/AEM.01574-07.

(13) Silvi, S.; Credi, A. Luminescent sensors based on quantum dot–molecule conjugates. Chemical Society Reviews 2015, 44 (13), 4275–4289, 10.1039/C4CS00400K. DOI: 10.1039/C4CS00400K.

(14) Lemon, C. M. Optical oxygen sensing with quantum dot conjugates. Pure and Applied Chemistry 2018, 90 (9), 1359–1377. DOI: doi:10.1515/pac-2018-0303 (accessed 2026-05-24).

(15) Otto, M. *Staphylococcus epidermidis*--the ’accidental’ pathogen. Nature reviews. Microbiology 2009, 7 (8), 555–567. DOI: 10.1038/nrmicro2182.

(16) Rusak, D. A.; James, W. H., III; Ferzola, M. J.; Stefanski, M. J. Investigation of Fluorescence Lifetime Quenching of Ru(bpy)32+ by Oxygen Using a Pulsed Light-Emitting Diode. Journal of Chemical Education 2006, 83 (12), 1857. DOI: 10.1021/ed083p1857.

(17) Ogawa, M.; Nakamura, T.; Mori, J.-i.; Kuroda, K. Luminescence of Tris(2,2‘-bipyridine)ruthenium(II) Cations ([Ru(bpy)3]2+) Adsorbed in Mesoporous Silica. The Journal of Physical Chemistry B 2000, 104 (35), 8554–8556. DOI: 10.1021/jp0015123.

(18) Kalyanasundaram, K. Photophysics, photochemistry and solar energy conversion with tris(bipyridyl)ruthenium(II) and its analogues. Coordination Chemistry Reviews 1982, 46, 159–244. DOI: 10.1016/0010-8545(82)85003-0.

(19) Corporation, N. U. HECZW Technical Specifications. 2024. https://nn-labs.com/collections/cadmium-based-quantum-dots/products/high-efficiency-water-soluble-cadmium-selenide-zinc-sulfide-cdse-zns-core-shell-quantum-dots-czw (accessed May 26, 2026).

(20) Jorge, P. A.; Maule, C.; Silva, A. J.; Benrashid, R.; Santos, J. L.; Farahi, F. Dual sensing of oxygen and temperature using quantum dots and a ruthenium complex. Anal Chim Acta 2008, 606 (2), 223–229. DOI: 10.1016/j.aca.2007.11.008.

(21) Christensen, G. D.; Baddour, L. M.; Simpson, W. A. Phenotypic variation of *Staphylococcus epidermidis* slime production *in vitro* and *in vivo*. Infection and Immunity 1987, 55 (12), 2870–2877. DOI: doi:10.1128/iai.55.12.2870-2877.1987.

(22) Goldstein, F. W.; Coutrot, A.; Sieffer, A.; Acar, J. F. Percentages and distributions of teicoplanin- and vancomycin-resistant strains among coagulase-negative staphylococci. Antimicrobial Agents and Chemotherapy 1990, 34 (5), 899–900. DOI: 10.1128/aac.34.5.899.

(23) Zonozi, R.; Wu, A.; Shin, J. I.; Secora, A.; Coresh, J.; Inker, L. A.; Chang, A. R.; Grams, M. E. Elevated Vancomycin Trough Levels in a Tertiary Health System: Frequency, Risk Factors, and Prognosis. Mayo Clin Proc 2019, 94 (1), 17–26. DOI: 10.1016/j.mayocp.2018.08.034.

(24) Chander, S.; Kumari, R.; Wang, H. Y.; Mohammed, Y. N.; Parkash, O.; Lohana, S.; Sorath, F. N. U.; Lohana, A. C.; Sadarat, F. N. U.; Shiwlani, S. Effect of low vs. high vancomycin trough level on the clinical outcomes of adult patients with sepsis or gram-positive bacterial infections: a systematic review and meta-analysis. BMC Infectious Diseases 2024, 24 (1), 1114. DOI: 10.1186/s12879-024-09927-4.

(25) Polonio, R. E.; Mermel, L. A.; Paquette, G. E.; Sperry, J. F. Eradication of biofilm-forming *Staphylococcus epidermidis* (RP62A) by a combination of sodium salicylate and vancomycin. Antimicrob Agents Chemother 2001, 45 (11), 3262–3266. DOI: 10.1128/aac.45.11.3262-3266.2001.

(26) Sakimura, T.; Kajiyama, S.; Adachi, S.; Chiba, K.; Yonekura, A.; Tomita, M.; Koseki, H.; Miyamoto, T.; Tsurumoto, T.; Osaki, M. Biofilm-Forming *Staphylococcus epidermidis* Expressing Vancomycin Resistance Early after Adhesion to a Metal Surface. BioMed Research International 2015, 2015 (1), 943056. DOI: 10.1155/2015/943056.

(27) Skovdal, S.; Hansen, L.; Ivarsen, D.; Zeng, G.; Büttner, H.; Rohde, H.; Jørgensen, N.; Meyer, R. Host factors abolish the need for polysaccharides and extracellular matrix-binding protein in *Staphylococcus epidermidis* biofilm formation. Journal of Medical Microbiology 2021, 70. DOI: 10.1099/jmm.0.001287.

(28) Dmitriev, R. I.; Papkovsky, D. B. Optical probes and techniques for O2 measurement in live cells and tissue. Cell Mol Life Sci 2012, 69 (12), 2025–2039. DOI: 10.1007/s00018-011-0914-0 .

(29) Stern, O.; Volmer, M. On the decay time of fluorescence. Phys. Z 1919, 20, 183–188.

(30) Jiang, B.; Ren, C.; Li, Y.; Lu, Y.; Li, W.; Wu, Y.; Gao, Y.; Ratcliffe, P. J.; Liu, H.; Zhang, C. Sodium sulfite is a potential hypoxia inducer that mimics hypoxic stress in *Caenorhabditis elegans*. JBIC Journal of Biological Inorganic Chemistry 2011, 16 (2), 267–274. DOI: 10.1007/s00775-010-0723-1.

(31) Heydorn, A.; Nielsen, A. T.; Hentzer, M.; Sternberg, C.; Givskov, M.; Ersbøll, B. K.; Molin, S. Quantification of biofilm structures by the novel computer program COMSTAT. Microbiology (Reading) 2000, 146 *( Pt* *10*), 2395–2407. DOI: 10.1099/00221287-146-10-2395 .

(32) Dalecki, A. G.; Crawford, C. L.; Wolschendorf, F. Targeting Biofilm Associated *Staphylococcus aureus* Using Resazurin Based Drug-susceptibility Assay. J Vis Exp 2016, (111). DOI: 10.3791/53925.

(33) Duniec, J. T.; Thorne, S. W. Environmental effects on fluorescence quantum efficiencies and lifetimes: a semiclassical approach. Journal of Physics C: Solid State Physics 1979, 12, 4109–4117.

(34) Ahmed, S.; Zang, Z.-W.; Yoo, K.; Ali, M. A.; Alfano, R. R. Effect of multiple light scattering and self-absorption on the fluorescence and excitation spectra of dyes in random media. Applied optics 1994, 33 13, 2746–2750.

(35) Hunter, R. C.; Beveridge, T. J. Application of a pH-sensitive fluoroprobe (C-SNARF-4) for pH microenvironment analysis in *Pseudomonas aeruginosa* biofilms. Applied and environmental microbiology 2005, 71 (5), 2501–2510. DOI: 10.1128/AEM.71.5.2501-2510.2005.

(36) Schlafer, S.; Raarup, M. K.; Meyer, R. L.; Sutherland, D. S.; Dige, I.; Nyengaard, J. R.; Nyvad, B. pH Landscapes in a Novel Five-Species Model of Early Dental Biofilm. PLOS ONE 2011, 6 (9), e25299. DOI: 10.1371/journal.pone.0025299.

(37) Schlafer, S.; Garcia, J. E.; Greve, M.; Raarup, M. K.; Nyvad, B.; Dige, I. Ratiometric imaging of extracellular pH in bacterial biofilms with C-SNARF-4. Appl Environ Microbiol 2015, 81 (4), 1267–1273. DOI: 10.1128/AEM.02831-14.

(38) Fulaz, S.; Hiebner, D.; Barros, C. H. N.; Devlin, H.; Vitale, S.; Quinn, L.; Casey, E. Ratiometric Imaging of the in Situ pH Distribution of Biofilms by Use of Fluorescent Mesoporous Silica Nanosensors. ACS Applied Materials & Interfaces 2019, 11 (36), 32679–32688. DOI: 10.1021/acsami.9b09978.

(39) Hao, L.; Li, J.; Kappler, A.; Obst, M. Mapping of heavy metal ion sorption to cell-extracellular polymeric substance-mineral aggregates by using metal-selective fluorescent probes and confocal laser scanning microscopy. Appl Environ Microbiol 2013, 79 (21), 6524–6534. DOI: 10.1128/aem.02454-13.

(40) Cotter, J. J.; O’Gara, J. P.; Casey, E. Rapid depletion of dissolved oxygen in 96-well microtiter plate *Staphylococcus epidermidis* biofilm assays promotes biofilm development and is influenced by inoculum cell concentration. Biotechnology and Bioengineering 2009, 103 (5), 1042–1047. DOI: 10.1002/bit.22335.

(41) Webster, C. M.; Shepherd, M. A mini-review: environmental and metabolic factors affecting aminoglycoside efficacy. World Journal of Microbiology and Biotechnology 2022, 39 (1), 7. DOI: 10.1007/s11274-022-03445-8.

(42) Pires, D. P.; Meneses, L.; Brandão, A. C.; Azeredo, J. An overview of the current state of phage therapy for the treatment of biofilm-related infections. Current opinion in virology 2022, 53, 101209.

(43) Strempel, N.; Strehmel, J.; Overhage, J. Potential application of antimicrobial peptides in the treatment of bacterial biofilm infections. Current pharmaceutical design 2014, 21 *1*, 67–84.

