## Supplementary information Figures S1-S2. for "*In situ* three-dimensional mapping of oxygen gradients in *Staphylococcus epidermidis* biofilms using a solution-based, ratiometric imaging platform"

**Supporting Information**

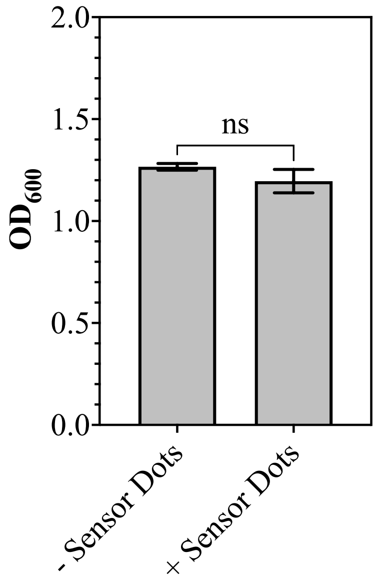


**Figure S1. PSt3 sensor dots do not alter biofilm growth.** *S. epidermidis* RP62A biofilms were grown for 24 hours with and without PSt3 oxygen sensor dots adhered to the interior of 8-well dishes (n=3). Biofilm biomass after 24-hours was quantified using $OD_{600}$ measurements and there was no statistically significant difference in biofilm growth with (+ sensor dots) and without (- sensor dots) PSt3 sensor dots indicating that the presence of PSt3 sensor dots does not alter overall biofilm growth. Statistical analysis was performed using an unpaired, two-tailed Welch's *t*-test (*ns:* not significant).

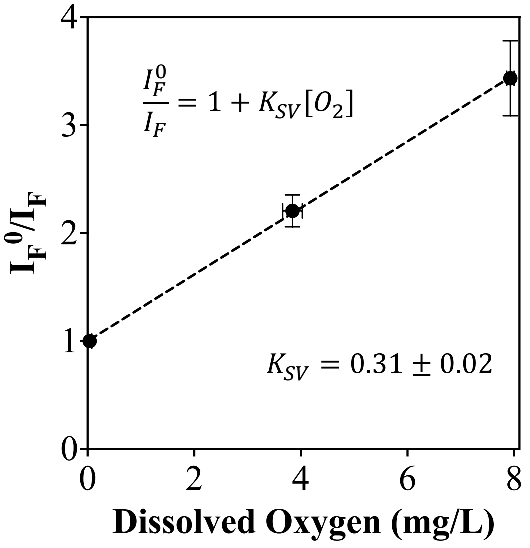


**Figure S2. Calibration of tris(2,2'-bipryridyl)dichlororuthenium(II) hexahydrate and oxygen insensitive CdSe/ZnS quantum dots in solution.** Ratiometric dissolved oxygen calibration curve for oxygen-sensing platform in solution with fluorescence intensity ratio ($I_F^0/I_F$) plotted against dissolved oxygen concentration (mg/L). The Stern-Volmer constant, $K_{SV}$, is 0.31 ± 0.02 L/mg for the calibration performed in solution, where the uncertainty represents the standard error of the fitted slope. Calibration points are for tryptic soy broth supplemented with 1 wt.% glucose ($TSB_g$) with dissolved oxygen concentrations of 0.04 ± 0.02 mg/L, 3.84 ± 0.09 mg/L, and 7.93 ± 0.03 mg/L as achieved using 4 g/L, 0.2 g/L, and 0 g/L sodium sulfite, respectively. The calibration curve indicates that there is a linear relationship between oxygen quenching and the fluorescence intensity ratio of tris(2,2'-bipryridyl)dichlororuthenium(II) hexahydrate to CdSe/ZnS quantum dots in solution from 0-8 mg/L dissolved oxygen.
